# pyFoldX: enabling biomolecular analysis and engineering along structural ensembles

**DOI:** 10.1101/2021.08.16.456210

**Authors:** Leandro G. Radusky, Luis Serrano

**Affiliations:** Centre for Genomic Regulation (CRG), The Barcelona Institute for Science and Technology, Barcelona 08003, Spain; Universitat Pompeu Fabra (UPF), Barcelona, Spain; ICREA, Pg. Lluis Companys 23, Barcelona 08010, Spain

## Abstract

Recent years have seen an increase in the number of structures available, not only for new proteins but also for the same protein crystallized with different molecules and proteins. While protein design software have proven to be successful in designing and modifying proteins, they can also be overly sensitive to small conformational differences between structures of the same protein. To cope with this, we introduce here pyFoldX, a python library that allows the integrative analysis of structures of the same protein using FoldX, an established forcefield and modeling software. The library offers new functionalities for handling different structures of the same protein, an improved molecular parametrization module, and an easy integration with the data analysis ecosystem of the python programming language.

**Availability and implementation:** pyFoldX is an open-source library that uses the FoldX software for energy calculations and modelling. The latter can be downloaded upon registration in http://foldxsuite.crg.eu/ and is free of charge for academics. Full details on installation, tutorials covering the library functionality, and the scripts used to generate the data and figures presented in this paper are available at https://github.com/leandroradusky/pyFoldX.

## Introduction

Outstanding advances in the experimentally determinable biomolecular space have been recently made (H. M. Berman, Vallat, and Lawson 2020); also, the last CASP competition (Senior et al. 2020) revealed striking improvements in the field of *in silico* biomolecular modelling. It is now increasingly likely that a protein of interest has several structures, alone or in complex with other biomolecules and ligands; however, protein design software users/researchers creating structural datasets commonly only select one of the available structures (normally the one with highest resolution and/or the highest sequence coverage) and disregard the rest. This loses the potential power of using different structures of the same protein and can result in significant differences in the analyses due to small conformational differences (Delgado Blanco et al. 2020) or the presence of different molecular partners along available structures (Kiel and Serrano 2014).

Thus, there is a need for bioinformatics tools that allow all available structures for a target protein to be considered when trying to diagnose mutational effects or to engineer new properties. These tools should take into account all structural information resulting from the formation of complexes and binding modes, as well as the distinct conformations arising from the crystallization conditions.

Here, we introduce pyFoldX, a python library powered by the FoldX suite (Delgado et al. 2019; Schymkowitz et al. 2005). pyFoldX enables full integration of standardized data analysis tools within the python programming language. The pyFoldX library mainly comprises two packages: i) the ***structure*** package, which contains classes to handle single structures or ensembles of structures **(Fig. 1A).** It is fully integrated with FoldX functionality (energy measurements, mutational modelling, etc.) and stores the results of energetic calculations in *pandas* data frames (McKinney 2015); and ii) the ***paramx*** package, which improves a functionality introduced in the latest version of FoldX, thus allowing for user-friendly parameterization of new molecules **(Fig. 1B)** based on standard atom types. Using these new functionalities, we created a comprehensive database of repaired PDB structures with minimized energy and improved structural quality. Our results from analyzing a comprehensive set of benign and pathogenic mutations mapped into protein structures underscored that considering the ensemble of available structures can improve the FoldX diagnosis. We have made the pyFoldX code open to the public and encourage the community to extend the library and/or request new specific features.

**Figure 1:**
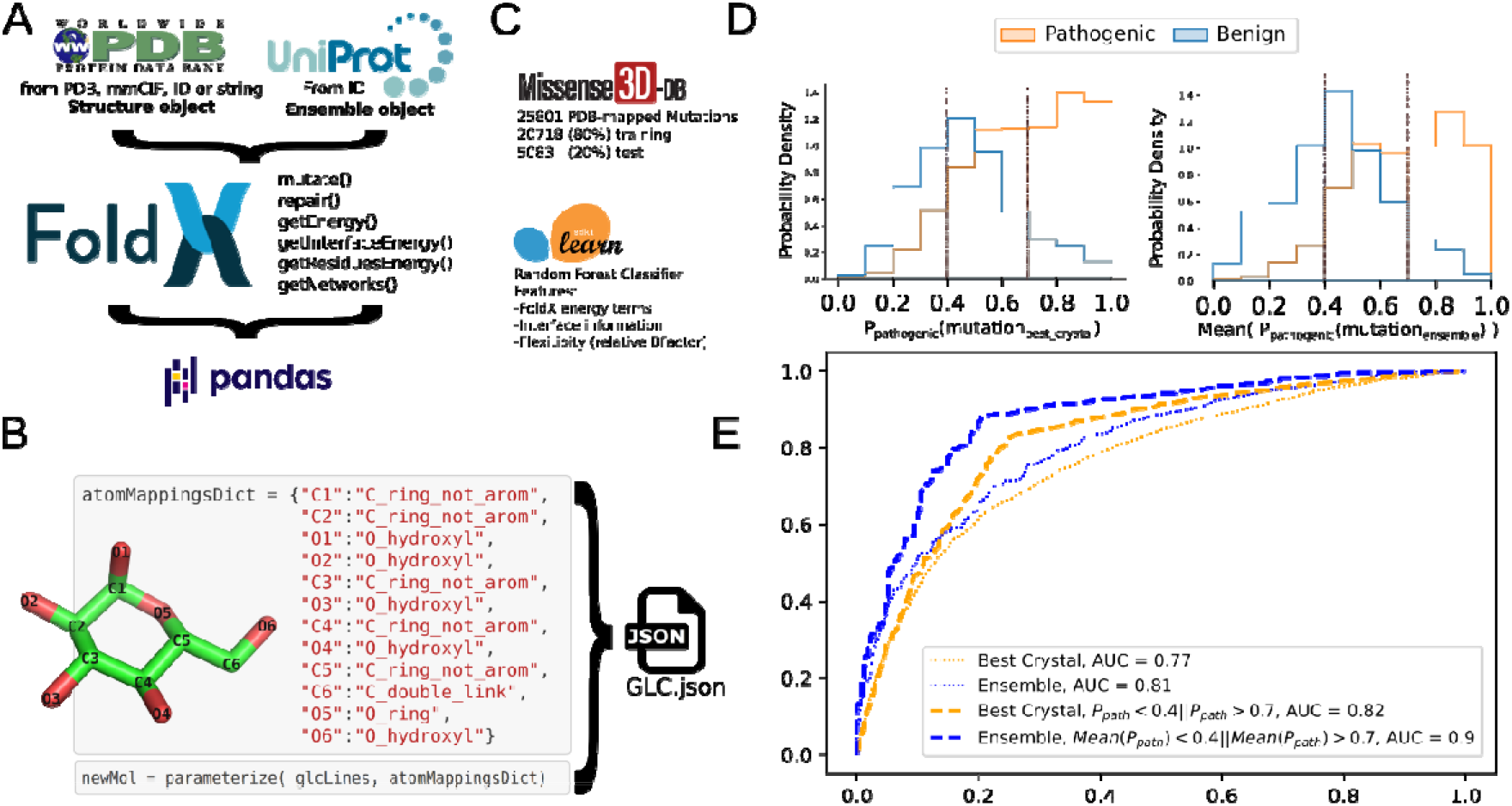
**(A)** pyFoldx structure-handling capabilities. Single structures can be instantiated from different formats, while ensembles of structures of the same protein can be instantiated from the protein’s Uniprot accession. FoldX commands can be executed into structures and ensembles, returning pandas dataframes with energies and, if applicable, objects with the transformed structures. (B) Example of parametrization of a glucose molecule with the pyFoldX paramx package. (C) Analyzed mutations data set description. To train a random forest classifier, 80% of the Missense3D-DB mutations were used in order to estimate the probability of belonging to the ‘pathogenic’ category. The remaining 20% were used for testing and analyzed by using the indicated structure in the database and the ensemble of good resolution structures for these proteins. (D) Histogram of probability of belonging to the ‘pathogenic’ category given by the created classifier for mutations mapped into their best structure by Missense3D-DB (left) and the mean of the probabilities for all crystals of good resolution along its ensemble (right). (E) ROC curve of mutation class prediction by the generated classifier taking into account best crystal (orange lines) or mean predictions for crystals along ensemble (blue lines). Thin lines: classifying mutations as pathogenic (P_pathogenic_>0.5) or benign (P_pathogenic_≤0.5). `Thick lines: mutations with no clear prediction are discarded` (0.4>P_path_>0.7). Overall, predictions are better when ensembles are considered and high accuracy is achieved (AUC = 0.9) when no clear predictions are discarded from the analysis.

## Functionality and Results

### Molecule Parameterization

A key feature that was introduced in the last version of FoldX was the possibility to parameterize small molecules not previously recognized by the software; this feature was effectively used the parameterization of RNA bases. We have now improved it in pyFoldX by allowing the auto parameterization of molecules through the simple definition of template atoms **(Table S1).** This feature will permit users to introduce new organic molecules in the FoldX force field using atoms that fit into the template atom set provided. A complete tutorial that demonstrates the parameterization of a glucose molecule complexed with a lectin is available online (Tutorials section of project’s github).

### Mutational Effect Diagnosis

Taking advantage of the integration of pyFoldX with *pandas*, we built a random forest classifier using the *sklearn* library (Trappenberg 2019). This allowed us to assess the accuracy of FoldX energetic features, along with other structural annotations, as well as to discriminate between benign and pathogenic mutations structurally mapped by the Missense3D-DB (Khanna et al. 2021). We trained the classifier with a random set encompassing 80% of the annotated mutations, with the remaining 20% used for testing **(Fig. 1C).** We then analyzed mutations along the ensemble of good resolution structures (resolution <2.0Å) for the same set of mutations. Based on the classifier probabilities for a mutation to belong to the ‘pathogenic’ or ‘benign’ category (P_pathogenic_=1-P_benign_), we found that mutations classified as benign tend to keep such classification along the ensemble of available structures for their proteins. This allowed us to achieve a better separation in the histogram of probabilities from both types of mutations **(Fig. 1D).** Notably, we observed a better discriminative power when we analyzed structural ensembles than when we used the best structure (e.g., in terms of resolution), giving us a highly accurate classification (AUC = 0.9) **(Fig. 1E).**

Note that the pyFoldX’s github webpage include: i) tutorials on how features that feed the classifier are computed; ii) the datasets of these features along single structures and ensembles mutations; and iii) scripts showing how the random forest classifier was created and used.

### FoldX-Repaired Protein DataBank

By combining the PDB_REDO (Joosten et al. 2014) structure refinement method with the Foldx *RepairPDB* command for sidechain energy minimization, we designed a pipeline using pyFoldX to refine the quality for virtually all the structures in the Protein Data Bank (H. Berman et al. 2007) **(Fig. S2).** We first examined the structures to determine if they contained organic compounds not recognized by FoldX; we then parameterized the most frequent ones. **Table S2** shows the list of new molecules now parameterized and recognized by FoldX.

In general, the resulting repaired structures presented lower energies when evaluated with independent force fields (Alford et al. 2017; Yang and Zhou 2008) **(Fig. S2A)** as well as improved structural quality features (as measured with WHAT_CHECK; (Dunbrack 2004) **(Fig. S2B).** If any of the independent force fields reported higher energies with respect to the original entries, the structures were further analyzed to determine potential flaws in FoldX modeling. The predicted high energies were due proline rotamers placed by FoldX but penalized by these force fields (Fig S3). FoldX uses a probabilistic method that depends on the previous phi, psi, and chi angles to generate rotamers; therefore, it is possible that some of the generated rotamers have a poor representation in the PDB structures. Note that all the generated structure files are available online for downloading or as object instantiation through pyFoldX structure package.

## Discussion and Perspectives

Here we present pyFoldX, a novel library fusing FoldX modeling functionalities within the python programming language. pyFoldX enables easy analyses of extensive structural datasets, and in-depth energy analyses and modelling along the ensemble of structures available for a single protein, in a user-friendly manner. Notably, comprehensive analysis using pyFoldX of mispredicted energy changes versus experimentally measured mutations should lead to improvements in the FoldX sidechain modeling routines and force field. Mutational effect diagnosis can now be improved by incorporating other non-structural features, and other machine learning strategies can be investigated to improve the built classifier. The possibility to parameterize novel organic molecules presents opportunities for ligand binding modelling; notably for carbohydrate molecules, given the ability of pyFoldX to handle mmCIF formatted files that correct historic errors on these kinds of interactions (Feng et al. 2021). Overall, we believe that this tool will provide the structural biology community with a more powerful way to access FoldX functionalities, useful for instance for determining effects of mutations or for engineering new properties into proteins/protein-ligand complexes.

## Acknowledgements

The authors would like to thank Javier Delgado Blanco for help on the molecule parameterization; Damiano Cianferoni for testing the package installation in different platforms; Veronica Raker for language editing; the Centre for Genomic Regulation (CRG) Technology & Business Development Office (TBDO) for support with licensing information; and the CRG Scientific Information Technologies (SIT) for distributed computing.

## Funding

This work was supported by funds from the European Regional Development Fund (ERDF) and the Generalitat de Catalunya (VEIS-001-P-001647, Programa Operatiu FEDER de Catalunya 2014-2020) and funds from Spanish Ministry of Science, Innovation and University (MCIU) (PGC2018-101271-B-I00). The authors also acknowledge the support of the Spanish Ministry of Science and Innovation to the EMBL partnership, the Centro de Excelencia Severo Ochoa and the Centres de Recerca de Catalunya (CERCA) Programme/Generalitat de Catalunya.

### Conflict of Interest

none declared.

## Notes

https://github.com/leandroradusky/pyfoldx

### Competing Interest Statement

The authors have declared no competing interest.

