## Supplementary Information for "pyFoldX: enabling biomolecular analysis and engineering along structural ensembles"

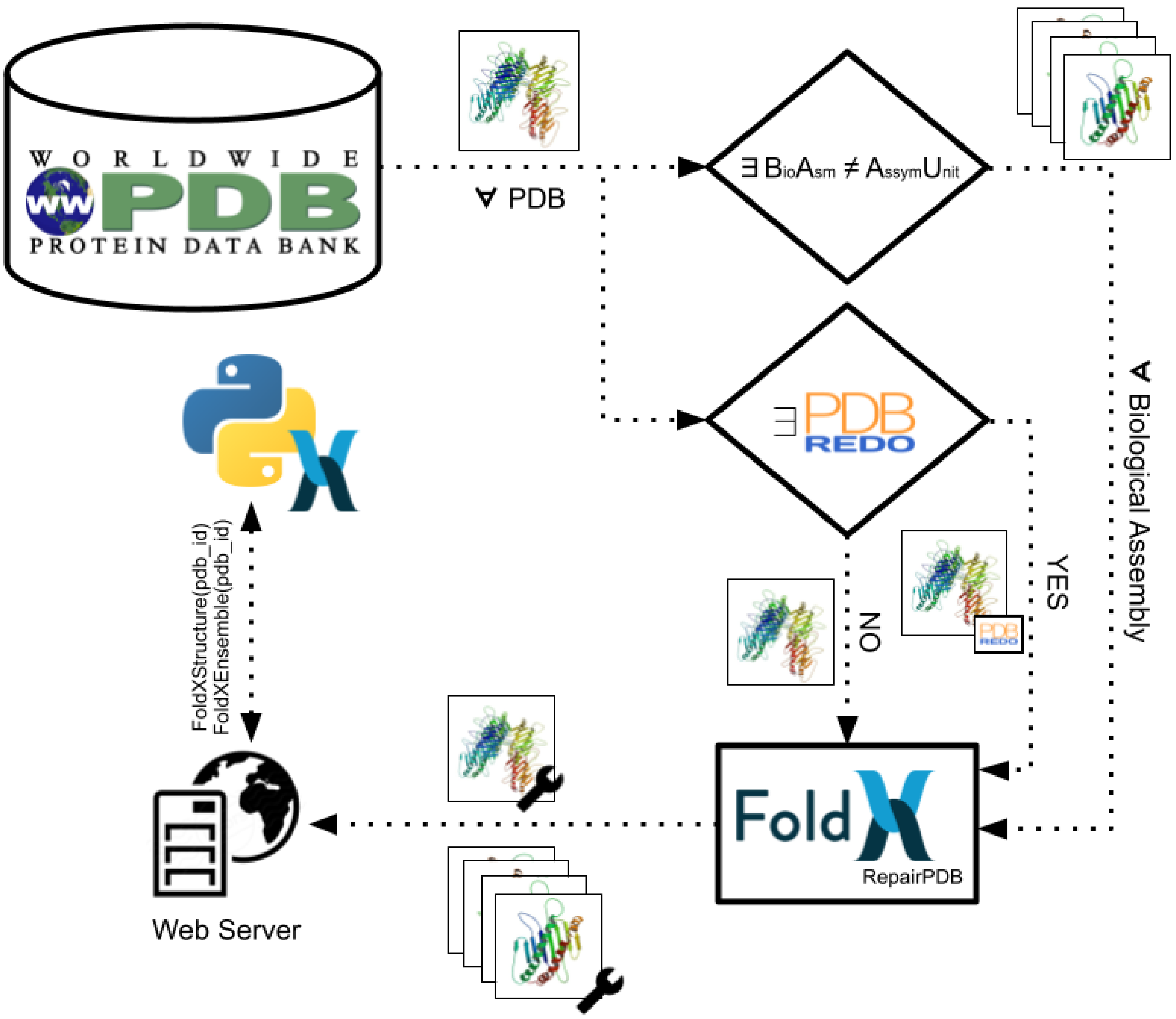


**Figure S1**: General pipeline describing the genesis of the Repaired PDB resource. When biological assemblies different of the asymmetric unit exists were repaired separately and stored to be accessible. The whole biological assembly was repaired taking the PDB_REDO entry when it exists, otherwise was taken as provided by the PDB. All the resulting structures are stored online and are accessible through the pyFoldX FoldXStructure and FoldXEnsemble class.

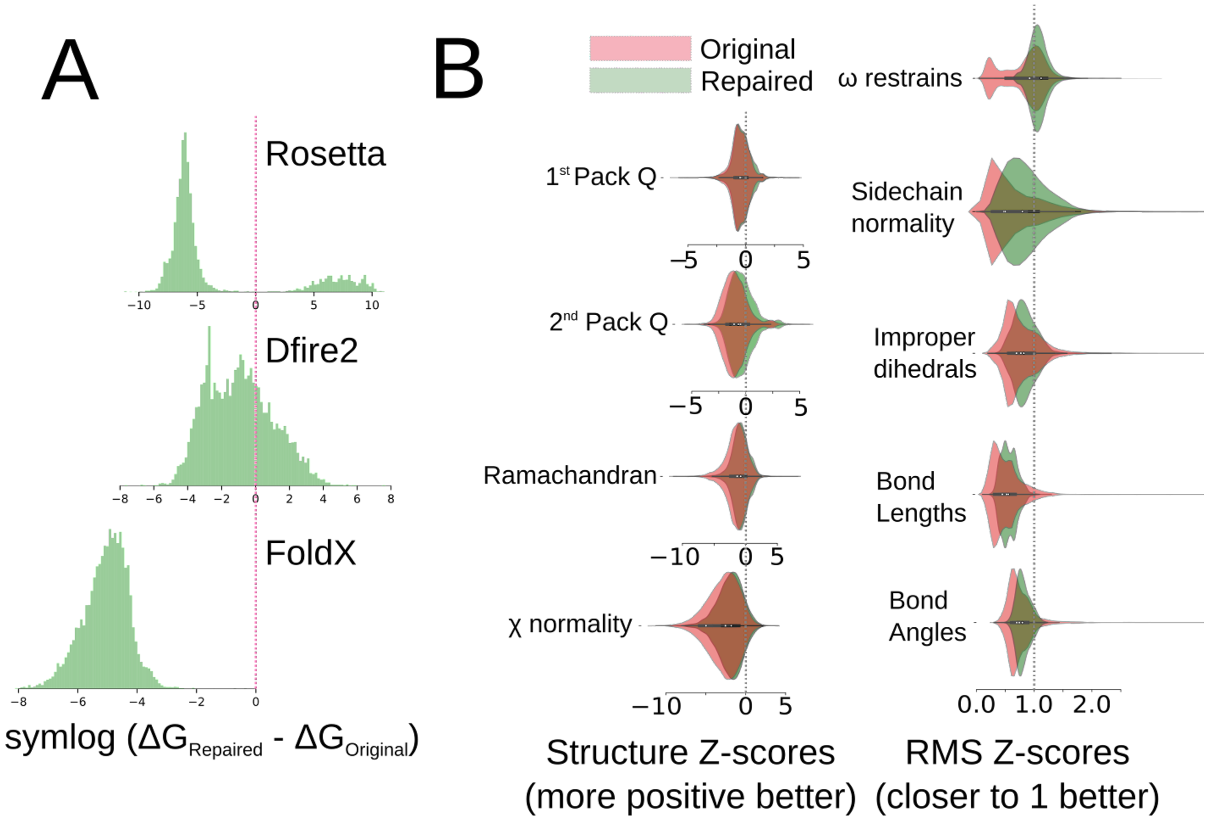


**Figure S2:** Evaluation of generated models. (A) Energetic evaluation presents tautologically better stabilities in FoldX forcefield and presents better energies for the majority of the models in the independent forcefields Rosetta(84%) and DFire2(76%). (B) Quality parameters measured with WHAT_CHECK ─violin plot histograms─ presents univocally better values for the generated models than for the original PDBs.

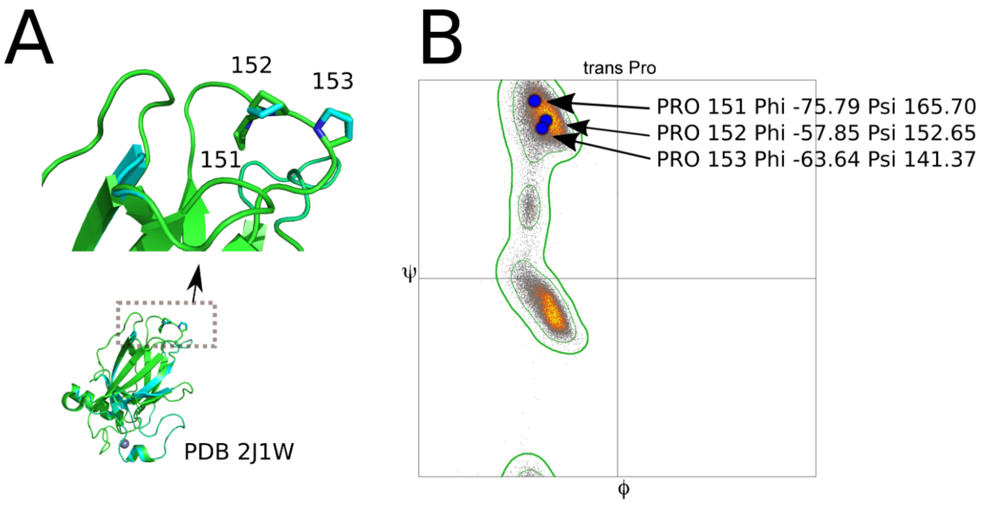


C

| Energy Term | ORIGINAL | REPAIRED | REPAIRED FIX PRO 151-153 |
|---|---|---|---|
| total_score | -223.15 | 224.262 | -231.11 |
| score | -223.15 | 224.262 | -231.11 |
| dslf_fa13 | 0 | 0 | 0 |
| fa_atr | -1092.351 | -1104.552 | -1104.552 |
| fa_dun | 356.179 | 536.937 | 355.289 |
| fa_elec | -268.221 | -284.466 | -284.466 |
| fa_intra_rep | 2.059 | 2.105 | 2.105 |
| fa_intra_sol_xover4 | 39.918 | 40.222 | 40.222 |
| fa_rep | 188.03 | 231.423 | 188.03 |
| fa_sol | 674.014 | 681.328 | 681.328 |
| hbond_bb_sc | -33.238 | -37.663 | -37.663 |
| hbond_lr_bb | -71.106 | -71.106 | -71.106 |
| hbond_sc | -25.014 | -22.483 | -22.483 |
| hbond_sr_bb | -25.892 | -25.892 | -25.892 |
| linear_chainbreak | 0 | 0 | 0 |
| lk_ball_wtd | -22.531 | -6.924 | -6.924 |
| omega | -0.937 | -0.937 | -0.937 |
| overlap_chainbreak | 0 | 0 | 0 |
| p_aa_pp | -37.121 | -37.121 | -37.121 |
| pro_close | 35.507 | 265.836 | 35.507 |
| rama_prepro | -4.264 | -4.264 | -4.264 |
| ref | 61.817 | 61.817 | 61.817 |
| time | 1 | 0 | 0 |
| yhh_planarity | 0 | 0 | 0 |

**Figure S3:** Example of penalized proline residues on repaired structures. **(A)** PDB entry 2J1W proline residues in positions 151, 152 and 153 were modified after FoldX Repair routine (cyan) respect to the original structure (green). **(B)** Repaired proline rotamers penalized by independent forcefields were checked in terms of psi and phi angles. **(C)** Rosetta forcefield energy terms for the original PDB entry, for the FoldX repaired structure and the Repaired structure with penalized prolines fixed (no repair). Despite the correctness of the repaired prolines, the fa_dun (full atom Dunbrack rotamer statistics) and pro_close (proline ring closeness) apply severe penalizations.

| Atom Name | Example atom | |
|---|---|---|
| O_hydroxyl | OG atom<br>SER molecule<br>(serine) | 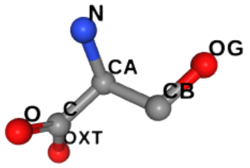 |
| O_ring | O4' atom<br>DA molecule<br>(adenosine) | 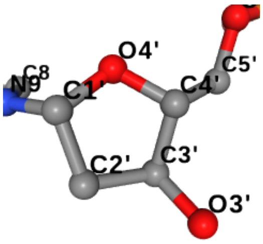 |
| O_minus | OD1 atom<br>ASP molecule<br>(aspartic acid) | 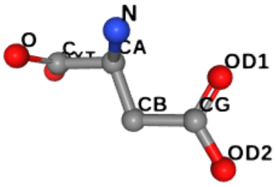 |
| O_carboxamide | OD1 atom<br>ASN molecule<br>(asparagine) | 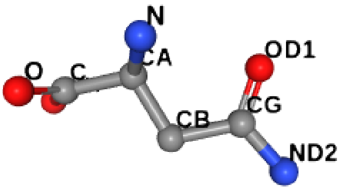 |
| N_amino | NZ atom<br>LYS molecule<br>(lysine) | 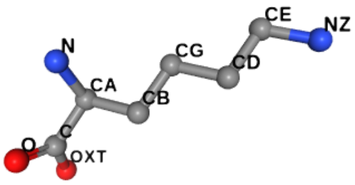 |
| N_guanidino | NH2 atom<br>ARG molecule<br>(arginine) | 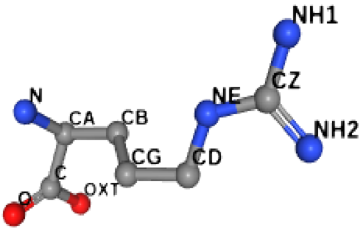 |

| | | |
|---|---|---|
| N_imidazol_plus | ND1 atom<br>HIS molecule<br>(histidine) | 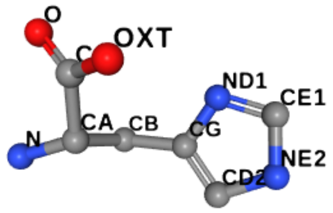 |
| N_imidazol_minus | NE2 atom<br>HIS molecule<br>(histidine) | 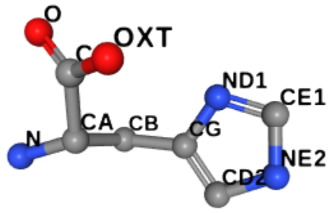 |
| N_pyrazole | N atom<br>PRO molecule<br>(proline) | 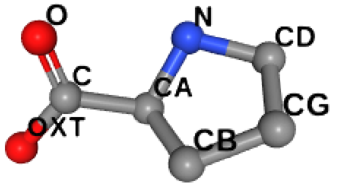 |
| N_amide | ND2 atom<br>ASP molecule<br>(asparagine) | 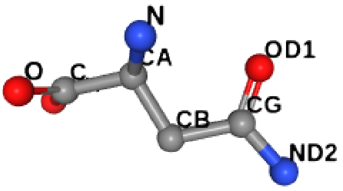 |
| C_ring_not_arom | CG atom<br>PRO molecule<br>(proline) | 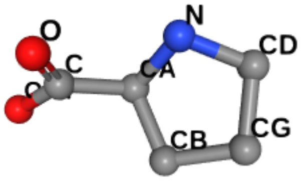 |
| C_ring_arom | CZ atom<br>PHE molecule<br>(phenilalanine) | 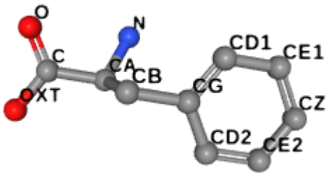 |

<table>
<tr><td>C_single_link</td><td>CG2 atom<br>THR molecule<br>(threonine)</td><td>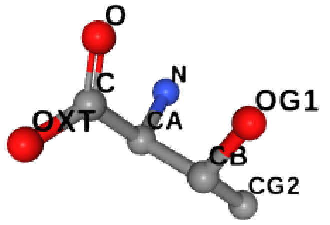
</td></tr>
<tr><td>C_double_link</td><td>CG atom<br>ARG molecule<br>(arginine)</td><td>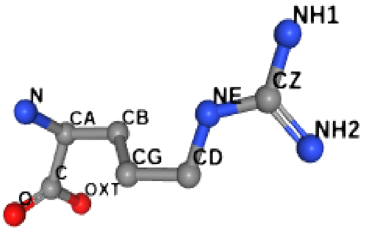
</td></tr>
<tr><td>C_triple_link</td><td>CG atom<br>LEU molecule<br>(leucine)</td><td>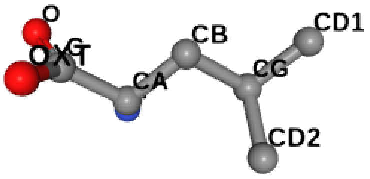
</td></tr>
</table>

**Table S1:** Template atoms that can be used in molecular parameterization with pyFoldX. Parameters will be copied from the indicated template atom to the specified atom in the new parameterized molecule conserving its chemical properties.

| PDB Code | Molecule Name | #Structs |
|---|---|---|
| **NAG** | 2-acetamido-2-deoxy-beta-D-glucopyranose | 6566 |
| **MAN** | alpha-D-mannopyranose | 3082 |
| **BMA** | beta-D-mannopyranose | 2945 |
| **GLC** | alpha-D-glucopyranose | 1507 |
| **FUC** | alpha-L-fucopyranose | 1432 |
| **GAL** | beta-D-galactopyranose | 1223 |
| **BGC** | beta-D-glucopyranose | 977 |
| **NDG** | 2-acetamido-2-deoxy-alpha-D-glucopyranose | 337 |

**Table S2:** Molecules previously unrecognized by FoldX parameterized using the pyFoldX's paramx module. From the molecules present in most structures (as free ligands) in the Protein Data Bank, those composed exclusively by atoms that can be mapped within the list of defined template atoms, and appearing in more than 100 structures, where parameterized. The parameters file for each compound in JSON format is available in the pyFoldX github webpage.
